# Engineering Bacteriophages to Overcome Complement-Mediated Neutralization

**DOI:** 10.64898/2026.11.10.765437

**Authors:** M. Abhyankar, P. Miernikiewicz, B. G. Rittmeyer, A. Khosravi, T. Luong, Z. Kaźmierczak, I Rybicka, A. Kłopot, P. L. Bollyky, K. Dąbrowska

**Author notes:** These authors contributed equally (M. Abhyankar, P. Miernikiewicz). These authors contributed equally (P. L. Bollyky, K. Dąbrowska). Corresponding Authors: Dr. Krystyna Dąbrowska Laboratory of Phage Molecular Biology, Hirszfeld Institute of Immunology and Experimental Therapy, Polish Academy of Sciences, Wrocław, Poland; Dr. Paul L. Bollyky, Division of Infectious Diseases and Geographic Medicine, Department of Medicine, Stanford University School of Medicine, Stanford, California, USA.

## Abstract

Bacteriophages (phages) are promising alternatives to antibiotics for treating antibiotic-resistant infections, but their clinical durability is limited by the host immune response against phages. Using phage T4 as a model and both C1q-KO and C3-deficient mice, we show that antibody recognition of phages triggers complement-dependent neutralization and clearance of phage particles *in vivo*. To mitigate this complement-mediated clearance, we engineer phages to display the complement-modulating Factor H binding peptide, a phage we call T4-FactorH.BP. Indeed, T4-FactorH.BP resists inactivation in human donor sera from individuals with pre-existing antibodies against T4 phage. Similarly, in mice immunized against T4 circulation of T4-FactorH.BP is prolonged *in vivo*. These results establish a critical role for the classical complement pathway in phage clearance and provide a path towards improving the pharmacokinetics of phage therapeutics *in vivo* by evading complement.

## INTRODUCTION

The growing global burden of multidrug-resistant (MDR) bacterial infections has revitalized interest in bacteriophage therapy, with numerous clinical successes reported over the past decade^1,2,3^. However, there are treatment failures as well^4,5^. We need to better understand the variables that influence phage efficacy *in vivo*.

Phage pharmacology is an important determinant of therapeutic outcomes, since efficacy depends not only on a phage’s ability to lyse its target bacterium, but also on whether it reaches the site of infection at sufficient concentration and for an adequate duration^6^. Intravenous administration, currently the predominant delivery route for systemic phage therapy, is limited by rapid clearance of phages from the bloodstream^7,8,9^. This clearance is largely mediated by the mononuclear phagocyte system (MPS) in the liver, spleen, and other tissues^6,10^, with substantial contributions from humoral factors, including antibodies^6,11^. The complement cascade is also implicated in phage inactivation^12–14^. However, the impact of complement on phage pharmacology *in vivo* and the extent to which each complement pathway contributes to these effects are unknown.

The complement system is a proteolytic cascade which contributes to the recognition, inactivation, and elimination of invading pathogens^15^. It can be activated via the classical pathway, typically initiated by antibodies bound to microbial antigens, the alternative pathway, which undergoes continuous low-level activation and provides an amplification loop, or the lectin pathway, when mannose-binding lectin recognize carbohydrate patterns on microbial surfaces^16^. Each pathway generates a C3 convertase that cleaves C3, the central component of the complement cascade, producing C3b that can covalently attach to microbial surfaces^17,18^. Although bacteriophages are not mammalian pathogens, there are indications that complement can reduce their infectivity^13,14,19^. Previous studies have demonstrated that heat-labile serum factors contribute to phage inactivation in the setting of antibodies^20,21^ but do not establish that this is due to the classical pathway or the alternative pathway. Further, it is unclear whether complement directly shapes phage pharmacokinetics *in vivo*.

To investigate this, we used bacteriophage T4, a well-characterized *Escherichia coli* phage with a myovirus morphotype characterized by a contractile tail, and T4 variants differing in the composition of their outer capsid proteins^20,22^. These included T4ΔHoc, a T4 mutant lacking functional Hoc (highly antigenic outer capsid protein)^23^, and T4ΔHocΔSoc, a T4 mutant lacking Hoc and Soc (small outer capsid protein)^23^. Hoc and Soc are nonessential outer capsid proteins displayed on the T4 head.

Here, using these phages and studies in complement-competent and complement-deficient mice, we demonstrate that complement directly influences phage pharmacokinetics. We also show phage neutralization by human complement *ex vivo*. Further, we report that engineering phages to display complement-modulating peptides on the phage capsid reduces complement mediated inactivation of phages *ex vivo* in human serum and improves phage pharmacokinetics *in vivo* in mice.

## RESULTS

### Phage capsid composition shapes complement-dependent pharmacokinetics *in vivo*

We first asked whether serum complement impacts phage pharmacokinetics. For these studies we used model phage T4^20^, as well as phage A3R, a *Staphylococcus aureus* phage with a myovirus morphotype that has been successfully used in phage therapy cocktails^24,25,26^ (**Figure 1A**).

**Figure 1.**
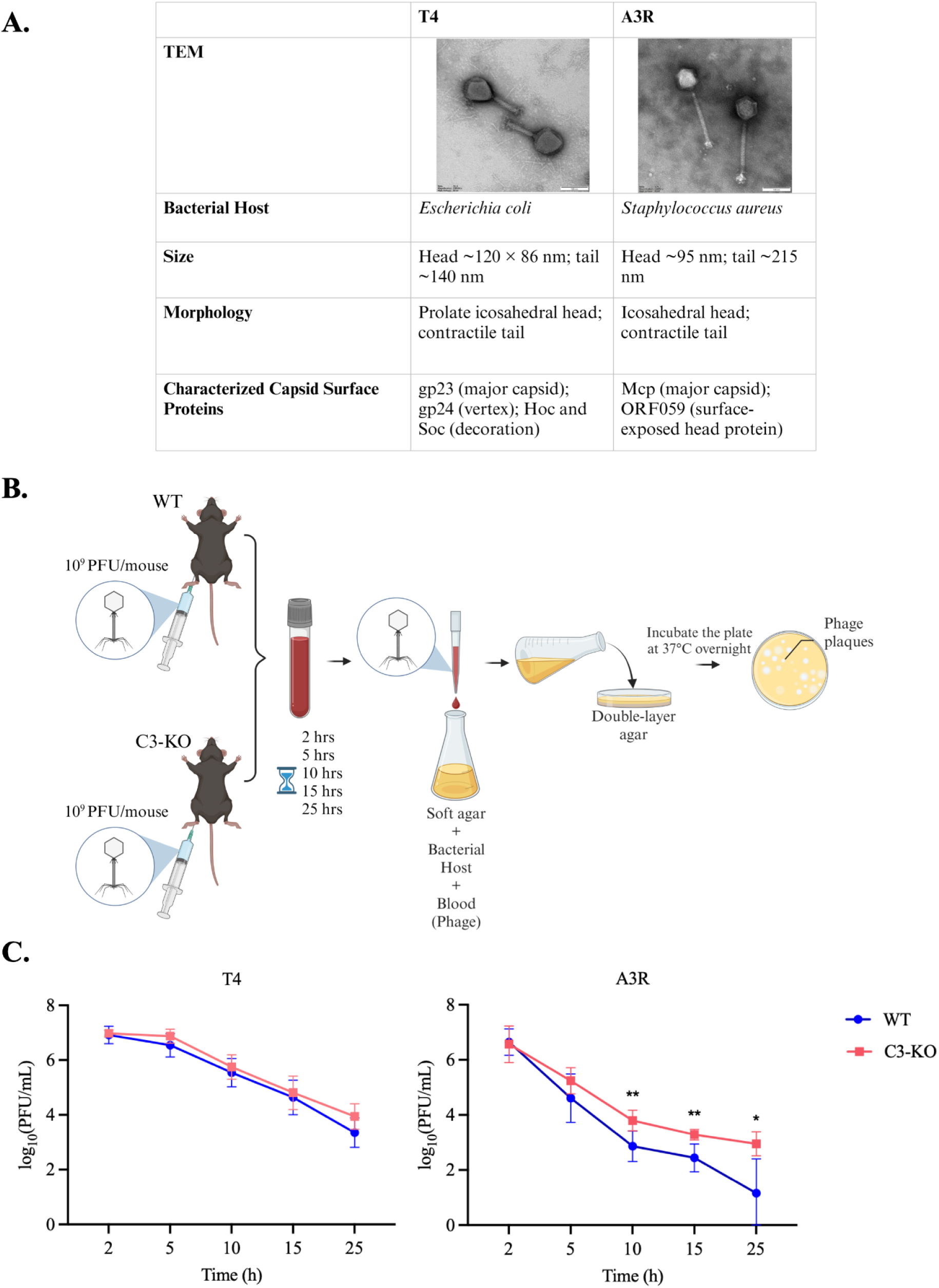
Susceptibility of phages A3R and T4 to serum complement neutralization. **A.** Characteristics of phages A3R^25^ and T4^22,27^. **B.** Schematic workflow for assessment of phage persistence *in vivo*. **C.** *In vivo* persistence of T4 and A3R in normal and C3-KO. WT and C3-KO were injected i.p. with 10^9^ plaque forming units (PFU)/mouse of T4 (N = 6) (A) or A3R (N = 8) phage preparations. Circulating infectious phage titers were measured at 2, 5, 10, 15, and 25 h following administration and expressed as log_10_(PFU/mL). Data are presented as mean ± SD. Statistical comparisons were performed using a mixed-effects model (REML) followed by Šídák’s multiple-comparisons test, comparing WT and C3-KO mice at each time point. No significant differences were detected for T4, whereas A3R titers were significantly higher in C3-KO mice at 10, 15, and 25 h. *Adjusted P < 0.05; **adjusted P < 0.01.

To assess the contribution of complement to phage persistence *in vivo*, we administered these phages to both wild-type (WT) and C3-deficient (C3-KO) mice. C3 is a central component of the cascade and C3 deficiency disrupts complement functions. Naïve mice lacking detectable phage-specific antibodies were used, and phage preparations were screened for lectin pathway activators, with none detected (data not shown). These conditions favoured assessment of alternative pathway activity. Phages were administered in a single dose intraperitoneally (IP), and phage titers in animal blood were monitored over the following 25 hours (**Figure 1B**). We observed that clearance of T4 phages was similar in naive WT and C3-KO mice and that titers declined over time in both strains (**Figure 1C**). In contrast, A3R titers were significantly higher in C3-KO than WT animals (**Figure 1C**). These findings indicate that complement contributes to the reduction of circulating active A3R phage, whereas no significant effect was detected for T4 under the conditions tested.

### Classical-pathway complement contributes to loss of circulating active T4 in pre-immunized mice

Because C3 deficiency did not significantly alter circulating T4 titers in naïve mice, we next asked whether complement contributes to T4 loss following induction of phage-specific antibodies.

To assess classical-pathway involvement, we compared T4 persistence in wild-type (WT) and C1q-deficient mice (C1q-KO), either naïve or pre-immunized with the T4 outer capsid protein Hoc (**Supplementary Figure. 2**). C1q is the recognition component of the C1 complex that initiates classical-pathway complement activation^28^. Mice received T4 intraperitoneally, and circulating infectious phage titers were measured over the following 25 hours.

**Figure 2.**
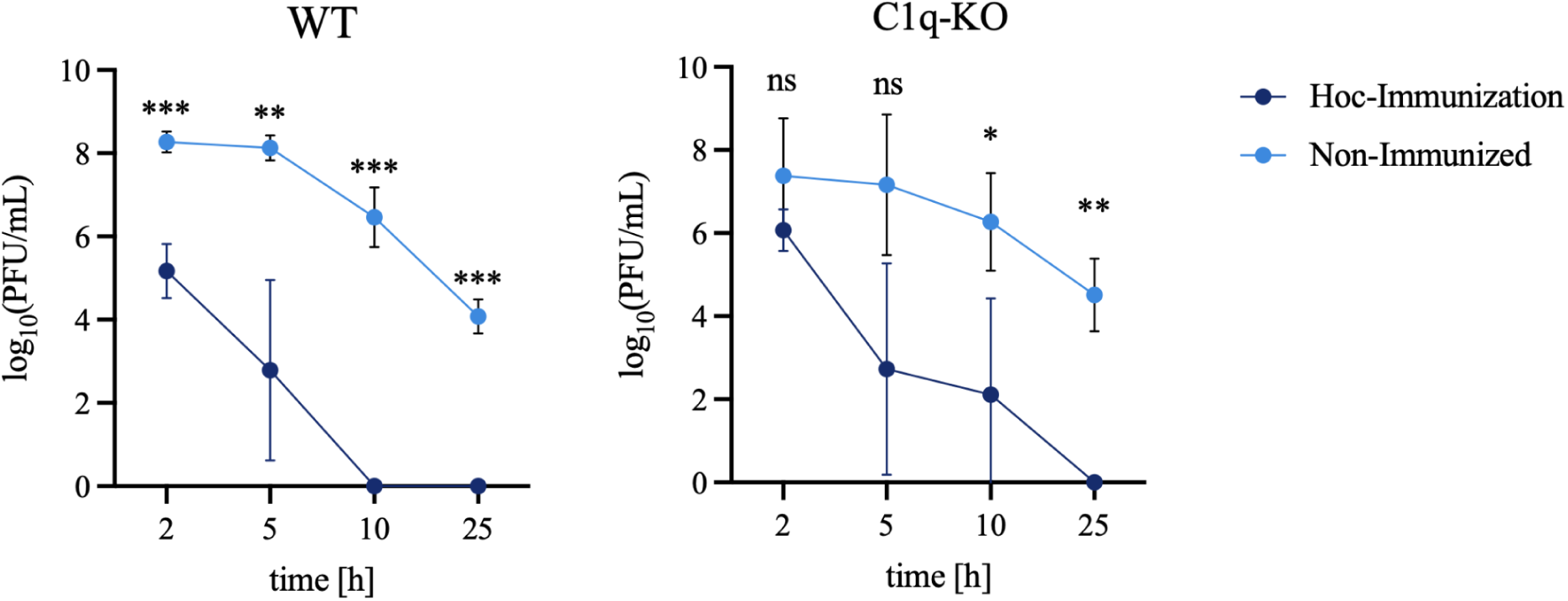
Inactivation of T4 phage in the blood of normal and C1q-KO mice. **A.** Circulating infectious T4 titers in WT mice following Hoc immunization compared with non-immunized controls. **B.** Circulating infectious T4 titers in C1q-KO mice following Hoc immunization compared with non-immunized controls. Mice were either non-immunized or pre-immunized with the T4 outer capsid protein Hoc 5 days before intraperitoneal (i.p.) administration of T4 phage (n = 6 mice per group initially). Blood was collected at 2, 5, 10, and 25 h following phage administration, and circulating infectious phage titers were determined by plaque assay. Data are expressed as log_10_(PFU/mL) and presented as mean ± SD. Statistical significance was determined using a mixed-effects model (REML) followed by Šídák’s multiple comparisons test comparing immunized and non-immunized mice at each time point (n = 5–6 observations per group per time point). ns, not significant; *P < 0.05, **P < 0.01, ***P < 0.001.

Following immunization with Hoc, phage-specific IgM antibodies were detected in both groups (**Supplementary Figure. 2**). In WT mice, immunization significantly reduced circulating infectious T4 titers compared with non-immunized controls at all measured time points (**Figure 2A**). In contrast, differences between immunized and non-immunized C1q-KO mice were not significant at 2 or 5 hours, although significant reductions emerged at 10 and 25 hours (**Figure 2B**).

Together, these findings demonstrate that Hoc immunization accelerates the loss of circulating infectious T4 in WT mice, while the early effects of immunization appear less pronounced in C1q-KO mice. This pattern suggests that classical-pathway complement contributes to antibody-associated phage clearance following immunization.

### Capsid protein composition shapes phage susceptibility to human complement neutralization *ex vivo*

Our *in vivo* findings implicated complement in phage clearance, prompting us to investigate whether human serum directly neutralizes phages and whether capsid protein composition influences their susceptibility. For these studies, we used phages A3R, T4, and T4ΔHoc variant. This engineered T4 variant lacks protein Hoc, a highly exposed T4 capsid protein that is dispensable for phage viability under laboratory conditions^29,30^. Hoc is present on WT T4 but absent from both its Hoc-deficient derivative T4ΔHoc^23^ and the therapeutic staphylococcal phage A3R.

Each phage was incubated with serum from healthy human donors containing phage-specific antibodies or with the same serum in which complement activity was disrupted by one of two independent approaches: heat treatment, to inactivate heat-labile complement proteins^31^, or treatment with cobra venom factor (CVF) which depletes complement through sustained activation and consumption of C3^32^.

We observed that all phages were neutralized by active human serum but that neutralization was prevented in inactivated serum. T4 was less susceptible to complement-mediated neutralization than A3R and T4ΔHoc (**Figure 3A**), supporting a protective role for Hoc against complement neutralization.

**Figure 3.**
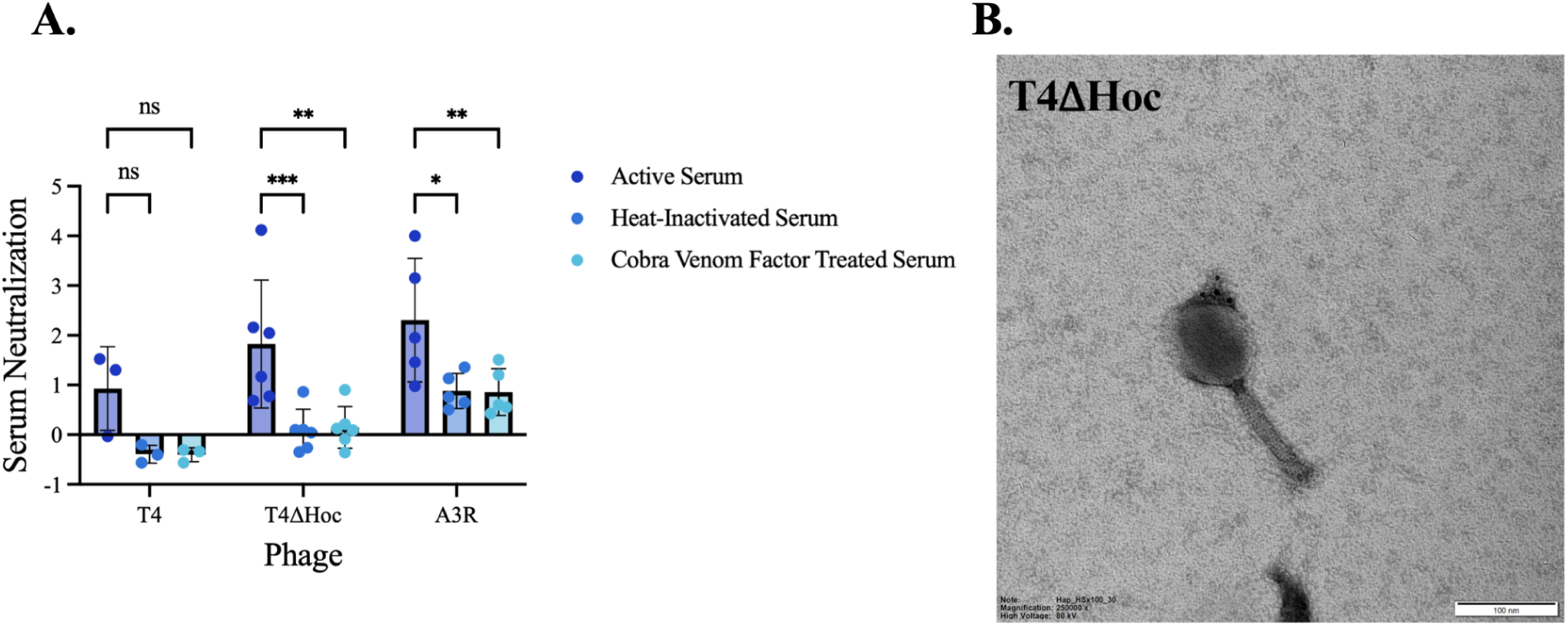
*Ex vivo* susceptibility of T4, T4ΔHoc, and A3R phages to human complement neutralization. **A.** Comparison of phage susceptibility to complement action *in vitro* in human serum. For these studies, untreated, heat-inactivated (1 h at 56°C), or CVF treated serum isolated from six healthy human volunteers were incubated 1:1 with phage preparations (10^7^ pfu/ml) for 1 h at 37°C. After incubation, phage activity was evaluated using full plate titering with the results expressed as log_10_(PBS_PFU/mL_/Serum Condition_PFU/ML_). **** adjusted p values < 0.0001 (2way ANOVA). ** adjusted p value = 0.0055 (2way ANOVA). **B.** Immunogold visualization of C3 deposition on T4ΔHoc. Scale bars: 100 nm

To further investigate complement interactions with the phage surface, C3 deposition on T4ΔHoc virions was examined by transmission electron microscopy (TEM) using immunogold labeling^33^. Following incubation with human serum, gold nanoparticles were readily detected in association with T4ΔHoc particles, with labeling predominantly localized to the capsid surface (**Figure 3B**). This observation was consistent with the susceptibility of T4ΔHoc to complement-mediated neutralization observed in the functional assay.

Together, these findings implicate complement C3 and antibodies against T4 in phage neutralization and support a role for Hoc in limiting C3 deposition on the T4 surface.

### Effects of bacteriophages on complement pathway activity

Having established that complement contributes to phage neutralization, we next asked whether bacteriophages themselves alter complement function. We assessed classical, lectin, and alternative complement pathway activity using the WIESLAB® Complement System assay. Neither T4 nor T4ΔHoc caused substantial changes in complement pathway activity under the conditions tested. No individual comparisons reached statistical significance following correction for multiple comparisons. T4ΔHoc was associated with a modest reduction in classical pathway activity, but this difference did not reach statistical significance (adjusted P = 0.0677; **Supplementary Figure. 1**).

### Engineering phages with short peptides to evade complement-mediated inactivation

Given our finding that human serum complement neutralizes T4ΔHoc phages, we then asked if T4ΔHoc phage can be protected from complement-mediated inactivation by specific molecular modifications of the phage surface involving peptides that prevent complement fixation.

We chose to target three different checkpoints of the pathway. A Factor H–binding peptide was selected to recruit the human complement regulator Factor H to the phage surface, thereby promoting the decay of the C3 convertase and limiting C3b amplification^34^. Compstatin, a well-characterized C3 inhibitor, was selected to block the cleavage of C3 into C3a and C3b. This mechanism inhibits downstream complement activation and C3b deposition on the phage surface, irrespective of the initiating pathway^35^. The third construct contained a hydrophilic peptide designed in this study as a potential antifouling agent. The peptide was rationally designed to enrich hydrophilic residues at the phage surface, with the goal of creating a highly hydrated interface while minimizing interactions that could promote complement deposition. This design was based on the established principle that highly hydrated surface chemistries can resist nonspecific protein adsorption by forming a hydration layer at the material–solution interface^36^. We hypothesized that displaying this peptide on the phage capsid would similarly reduce nonspecific complement protein adsorption without directly targeting a specific complement component.

We engineered these three peptides (**Figure 4A**) for capsid display by fusion to Soc, a second non-essential outer capsid protein present on phage T4. T4ΔHocΔSoc, which lacks both Hoc and Soc, was used as the platform accepting peptide-Soc fusion proteins (**Figure 4B, C**). Following incorporation, the engineered phages were Soc⁺, but Soc also carried the displayed peptides. T4ΔHoc, a T4 variant lacking Hoc but retaining native Soc, was used as a control, providing the corresponding Hoc⁻/Soc⁺ capsid background without peptide modification (**Figure 4B**). Phages displaying compstatin, a factor H-binding peptide, or a hydrophilic peptide were designated T4-Comp, T4-FactorH.BP, and T4-HydroP, respectively. Together, these engineered phages were designed for testing three complementary strategies: inhibition of C3 cleavage by T4-Comp, recruiting mammalian factor down-regulating C3 activation by T4-FactorH.BP, and reduction of complement interactions with the phage surface by T4-HydroP.

**Figure 4.**
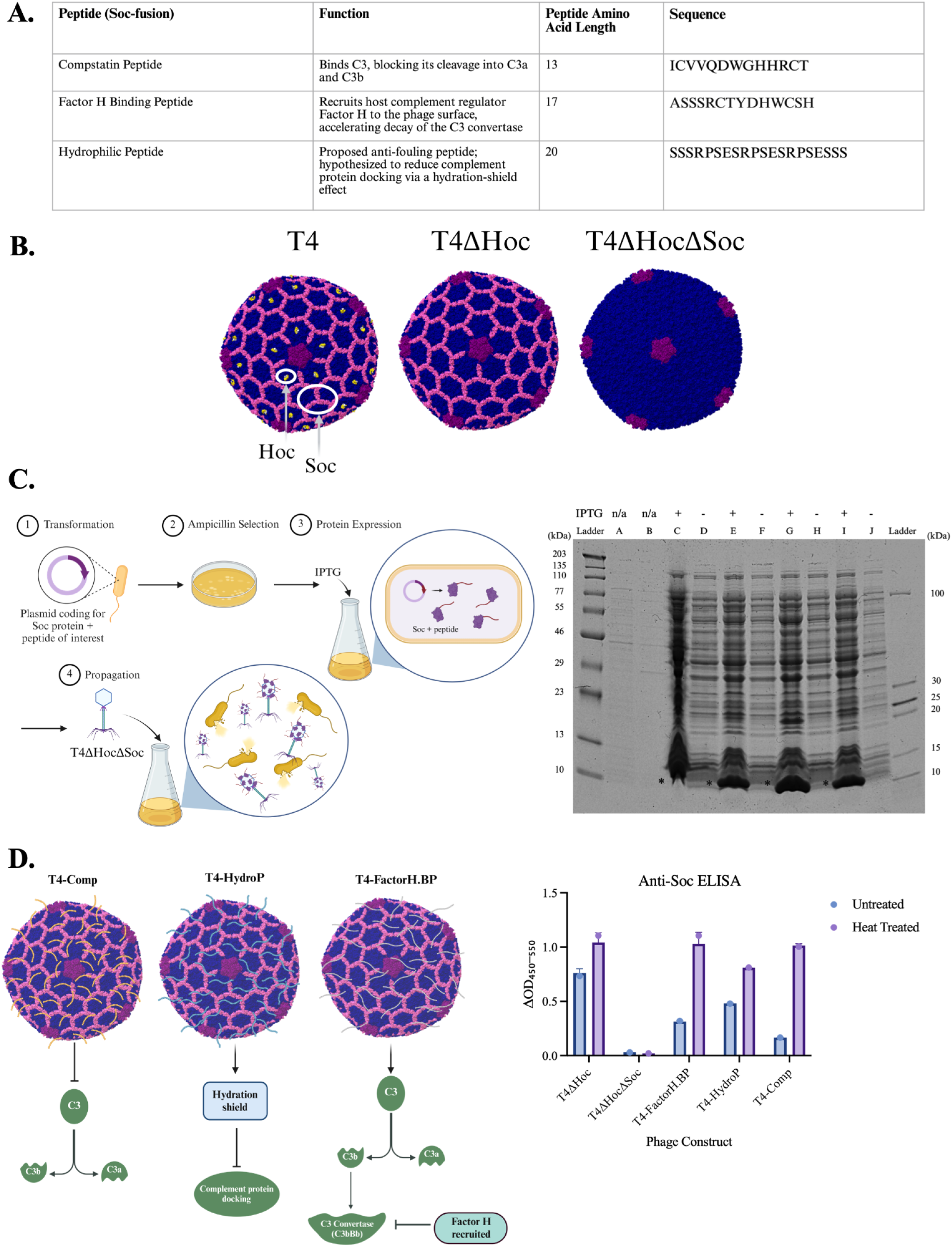
Engineering phages to evade complement-mediated inactivation. **A**. Peptides with anti-complement activity **B.** Structure-based representations of the T4, T4ΔHoc, and T4ΔHocΔSoc capsids generated in UCSF ChimeraX from the T4 isometric capsid structure (PDB: 5VF3; EMDB: EMD-8661). T4ΔHoc and T4ΔHocΔSoc were represented by removal of Hoc and Hoc/Soc, respectively, from the parental structural model to illustrate their outer capsid protein composition. **C.** Plasmid-mediated phage display by trans complementation: the capsid protein fusion was supplied in trans by the host bacterium and incorporated into progeny phage particles during assembly, without modification of the phage genome. SDS-PAGE analysis of *E. coli* lysates before and after IPTG induction, showing increased expression of Soc–peptide fusion proteins (∼10 kDa) after induction. **D.** Expression and capsid incorporation of peptide–Soc fusion proteins. Anti-Soc ELISA assessing incorporation of Soc–peptide fusion proteins onto engineered phage capsids, with and without heat treatment (80°C, 30 min). Heat treatment was applied to increase the accessibility of Soc proteins, which may otherwise be partially shielded by the intact phage capsid. T4ΔHoc and Soc-deficient T4ΔHocΔSoc served as positive and negative controls, respectively. Absorbance is reported as A_450-550_.

Expression of the peptide–Soc fusion proteins in host bacteria was confirmed by SDS-PAGE (**Figure 4C**). Incorporation of the peptide–Soc fusions into the phage capsid was conducted in the fusion-expressing bacteria infected with T4ΔHocΔSoc. Fusion incorporation was verified by anti-Soc ELISA using isolated and purified phages T4-Comp, T4-FactorH.BP, and T4-HydroP with controls: T4ΔHoc (Soc-positive) and T4ΔHocΔSoc (Soc-negative). All three fusion phages produced stronger ELISA signals than T4ΔHocΔSoc, particularly after heat treatment that enhances exposition of capsid components, confirming capsid incorporation of the peptide–Soc fusions (**Figure 4D**).

### Capsid-displayed anti-complement peptides protect phages *ex vivo* in human serum

To determine whether capsid-displayed complement inhibitors protect phages from serum-mediated neutralization, we compared three engineered phages: T4-Comp, T4-FactorH.BP, and T4-HydroP, with the parental T4ΔHoc phage in sera from three independent donors who previously exhibited neutralizing effects targeting T4ΔHoc (**Figure 3A**).

We observed that Factor H recruitment and C3 inhibition by Compstatin consistently protected phages across all tested sera, while the effect of hydrophilic surface modification was weaker and donor-dependent (**Figure 5**). These findings demonstrate that capsid-displayed peptides can reduce phage susceptibility to human serum complement.

**Figure 5.**
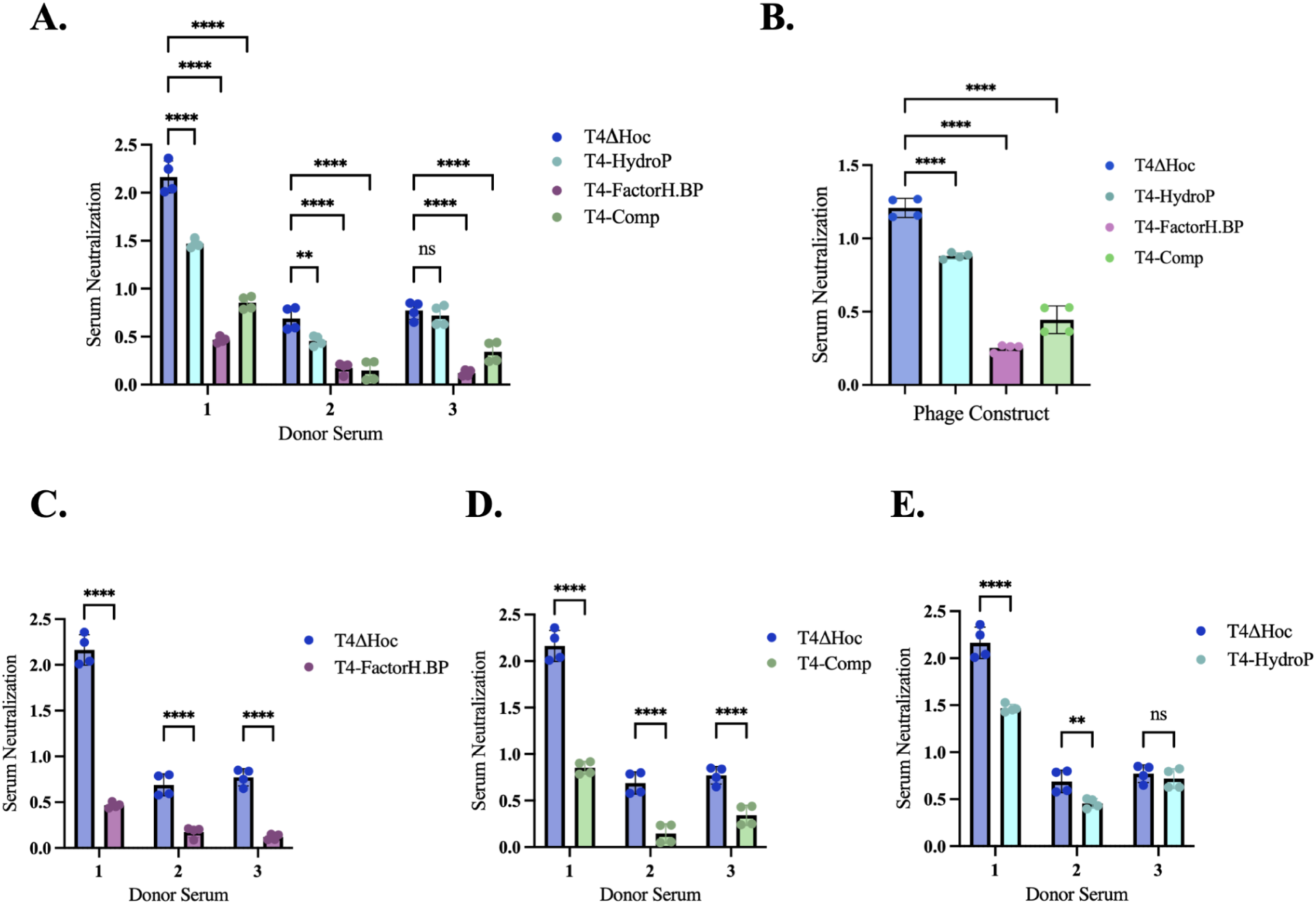
Capsid-displayed anti-complement peptides protect phages in human serum. **A** *Ex vivo* serum neutralization of T4ΔHoc and peptide-displaying phages (T4-HydroP, T4-FactorH.BP, and T4-Comp) in active human serum from three independent donors. **B** Summary of serum neutralization averaged across the three donors. **C–E.** Individual donor serum comparisons of T4ΔHoc with T4-FactorH.BP (**C**), T4-Comp (**D**), and T4-HydroP (**E**). Phages were incubated 1:1 with active human serum for 1 h at 37°C, and titers were determined by plaque assay. Serum-mediated neutralization was calculated as log_10_(PBS_PFU/mL_/Serum Condition_PFU/ML_). Points in (**A, C–E**) represent replicate measurements (n = 4 per condition), whereas points in (**B**) represent replicate values averaged across the three donors (n = 4 per construct). Bars represent mean ± SD. Statistical comparisons in (**A, C–E)** were performed using two-way ANOVA followed by Dunnett’s multiple-comparisons test, comparing each peptide-displaying phage with T4ΔHoc within each donor serum. Statistical comparisons in (**B**) were performed using ordinary one-way ANOVA followed by Dunnett’s multiple-comparisons test, comparing each peptide-displaying phage with T4ΔHoc (all adjusted P < 0.0001). **Adjusted P < 0.01; ****adjusted P < 0.0001; ns, not significant.

### Capsid-displayed peptides improve phage persistence *in vivo*

To determine whether anti-complement peptides also improved phage persistence *in vivo*, we compared circulating titers of T4-HydroP and T4-FactorH.BP with the control phages T4ΔHoc and T4ΔHocΔSoc in mice. T4-Comp was not evaluated in this model because compstatin exhibits species-specific activity toward primate C3 and does not inhibit murine C3^37^.

Since phage-specific antibodies contributed substantially to phage neutralization by serum complement (**Figure 2**), we pre-immunized mice with the T4 major capsid protein (gp23) to induce phage-specific antibodies that do not target the tail and fiber proteins involved in infection, thereby avoiding direct antibody-mediated phage neutralization. Efficient induction of phage-specific antibodies was confirmed by ELISA (**Supplementary Figure 3**).

Fourteen days following immunization, phages were administered intraperitoneally, and circulating infectious phage titers were monitored over 6 hours (**Figure 6A**). Titers declined over time across all groups. Both T4-FactorH.BP and T4-HydroP maintained significantly higher circulating titers than T4ΔHoc at 1, 3, and 6 h (**Figure 6B,C**). Compared with T4ΔHocΔSoc, T4-FactorH.BP also maintained significantly higher titers at all three time points (**Figure 6D**), whereas T4-HydroP showed a significant improvement only at 6 h (**Figure 6E**). No significant differences were detected between the control phages T4ΔHoc and T4ΔHocΔSoc at any time point (**Figure 6F**). No significant differences were detected between the two peptide-displaying phages at any time point (data not shown). These findings indicate that capsid display of the Factor H-binding and hydrophilic peptides improves phage persistence in circulation *in vivo*.

**Figure 6.**
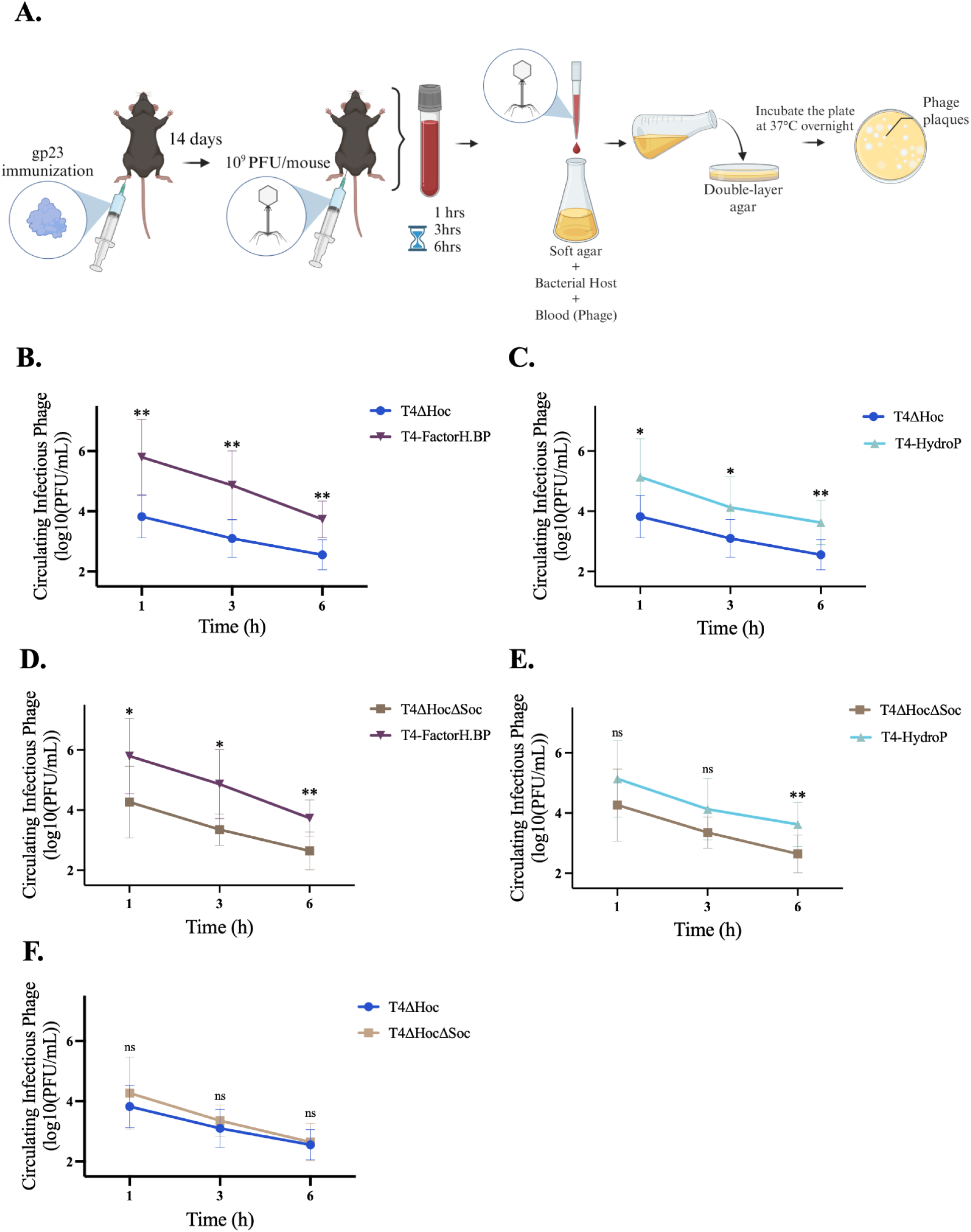
Anti-complement peptides improve phage persistence in *in vivo* murine models. (A) Schematic of the experimental design. Mice were immunized with gp23 14 days before intraperitoneal (i.p.) administration of 1 × 10^9^ PFU/mouse of T4ΔHoc (n = 11), T4ΔHocΔSoc (n = 13), T4-HydroP (n = 13), or T4-FactorH.BP (n = 10). Blood was collected at 1, 3, and 6 h after phage administration, and circulating phage titers were determined by plaque assay. (B–C) *In vivo* pharmacokinetics of T4-FactorH.BP (B) and T4-HydroP (C) compared with T4ΔHoc control phage. (D–E) *In vivo* pharmacokinetics of T4-FactorH.BP (D) and T4-HydroP (E) compared with T4ΔHocΔSoc control phage. (F) Comparison of T4ΔHoc and T4ΔHocΔSoc pharmacokinetics. Circulating phage titers are expressed as log_10_(PFU/mL), with data presented as mean ± SD. Statistical comparisons were performed using mixed-effects models (REML), with repeated measurements matched within individual mice, followed by Šídák’s multiple-comparisons test comparing phage constructs at each time point. *Adjusted P < 0.05; **adjusted P < 0.01; ns, not significant.

## DISCUSSION

The success of phage therapy depends on whether a phage can kill its bacterial target, but also on whether it remains bioactive within the mammalian host long enough to do so.

Here, we identify complement as a determinant of phage persistence in circulation and show that its effects depend on both the phage surface and the immune state of the host. Importantly, we further show that this interaction is not fixed: engineering the phage capsid with complement-modulating peptides can reduce serum neutralization and prolong the persistence of infectious phage *in vivo*, providing a strategy for improving phage bioavailability.

Our findings implicate the classical complement pathway in T4 phage inactivation in serum. Following immunization against the T4 capsid protein Hoc, infectious T4 persisted longer in C1q-deficient mice than in wild-type animals. C1q provides the recognition arm of the C1 complex that initiates classical-pathway activation^28^, making these observations consistent with an increased contribution of antibody-dependent classical complement activity after immunization. This distinction may be particularly relevant to phage therapy regimens when repeated phage administration is required, as each exposure has the potential to alter the immune response encountered by subsequent doses. Anti-phage antibodies may also be present in people prior to phage therapy^20,25,38^. Our findings suggest that the pharmacokinetic behavior of a phage cannot necessarily be inferred from its behavior in an immunologically naïve host.

Although complement contributes to phage neutralization, our results suggest that these interactions do not necessarily compromise complement function. Neither T4 nor T4ΔHoc substantially reduced classical, lectin, or alternative pathway activity under the conditions tested. The modest reduction in classical pathway activity observed with T4ΔHoc, although not statistically significant following correction, could reflect complement activation and consumption during incubation with a high phage concentration (10^10^ PFU/mL), rather than direct inhibition. Importantly, these in vitro conditions may not reflect those encountered *in vivo*, where complement components are continuously replenished. Thus, although bacteriophages are susceptible to complement-mediated neutralization, their interactions with complement do not appear to substantially impair overall pathway activity under the conditions examined.

Our comparison of T4 and A3R suggests that the role of complement in phage clearance is phage species-specific. C3 deficiency altered the persistence of A3R but had no detectable effect on T4 in naïve animals. This distinction is important because phages present extraordinarily diverse surfaces to the mammalian immune system. Capsid architecture, decoration proteins, surface chemistry, and associated bacterial products all vary among phages and may influence complement recognition.

The heterogeneity observed in healthy human sera reinforces the principle that complement-dependent neutralization differs substantially among donors. The degree of protection afforded by the engineered constructs was likewise donor dependent. Individual variation in pre-existing antibody repertoires and complement activity could therefore contribute to the variable pharmacokinetics reported for therapeutic phages^6^. Although the present study was not designed to identify the determinants of this variation, these observations raise the possibility that phage pharmacokinetics reflects an interaction between properties of the therapeutic particle and the immune history of the recipient. Establishing whether phage-specific antibody titers, complement activity or other serum features predict neutralization will require larger donor cohorts in which these variables are measured directly.

Our experiments with the T4 surface further suggest that capsid composition contributes to complement susceptibility. Hoc is a highly exposed, non-essential outer capsid protein^23,29^, and comparison of T4 with its Hoc-deficient derivative T4ΔHoc revealed altered susceptibility to human serum together with detectable C3 associated with T4ΔHoc particles by immunogold microscopy. These observations are consistent with the idea that decoration proteins influence how complement encounters the virion surface. They do not, however, establish whether Hoc directly prevents C3 deposition since removal of Hoc may alter several properties of the capsid simultaneously.

These findings also raise the possibility of engineering the phage surface to alter its interactions with the immune system. We used the T4 Soc display system to alter the phage surface through three distinct strategies: inhibition of C3 with Compstatin, recruitment of the endogenous complement regulator Factor H and presentation of a hydrophilic peptide intended to reduce nonspecific protein interactions. Despite their different designs, capsid-displayed peptides reduced serum neutralization, and the Factor H-binding and hydrophilic constructs improved persistence of infectious phage in pre-immunized mice. Together, these results provide proof of principle that complement susceptibility can be modified at the level of the therapeutic particle itself.

There are some limitations to the study. Although SDS–PAGE and anti-Soc ELISA confirmed peptide–Soc expression and incorporation into phage preparations, they did not quantify display occupancy, limiting our ability to relate peptide density to complement protection. In addition, circulating PFU provides a functional but incomplete measure of phage pharmacokinetics, as loss of infectious phage from blood may reflect neutralization, cellular uptake, tissue sequestration, or redistribution. Future studies quantifying biodistribution will be needed to define the mechanisms underlying the observed protection.

In summary, our findings suggest that mammalian immune recognition and the classical complement pathway in particular should be considered alongside bacterial host range, lytic activity and resistance evolution when optimizing therapeutic phages. Moreover, this work provides a framework for engineering phages with improved compatibility with host immunity and more predictable pharmacokinetics, with the potential to improve phage therapy efficacy.

## MATERIALS & METHODS

### Ethics statements

All animal experiments were performed according to EU directive 2010/63/EU for animal experiments and were approved by the 1st Local Committee for Experiments with the Use of Laboratory Animals, Wrocław, Poland (project no. 06/2017, 66/2017, 44/2018/P1). Blood samples from human volunteers were collected with their written consent, and in accordance with the local Commission of Bioethics, Regional Specialized Hospital in Wroclaw (approval no. KB/nr2/rok2017.

### Human serum collection and preparation

Human serum samples were obtained from healthy individual donors under Stanford IRB IRB-28425 via venous blood draw collected into BD Vacutainer™ Plastic Serum Tubes (silicone-coated, conventional stopper). Blood was allowed to clot at room temperature, after which serum was separated and clarified by centrifugation twice at 2,200 rpm for 10 min. Serum was aliquoted and stored at −80°C until use; samples were thawed immediately prior to experiments and were not subjected to repeated freeze–thaw cycles.

### Bacteriophage propagation

T4 bacteriophage was obtained from the American Type Culture Collection (ATCC), whereas T4ΔHoc and A3R were obtained from the Phage Collection of the Hirszfeld Institute of Immunology and Experimental Therapy (HIIET). Phages were propagated on their respective bacterial hosts, *Escherichia coli* B (ATCC; T4 and T4ΔHoc) and *Staphylococcus aureus* (HIIET Phage Collection; A3R), and purified as previously described^39^. Briefly, phage lysates were clarified by centrifugation, filtered through 0.22-µm membranes, and subjected to tangential flow filtration using a 750-kDa molecular weight cutoff hollow-fiber cartridge for buffer exchange into PBS and concentration. All phage preparations were additionally purified by endotoxin-affinity chromatography using EndoTrap HD (Lionex GmbH, Germany), dialyzed against PBS using membranes with a molecular weight cutoff of 1,000 kDa, and finally filtered through 0.22-µm membranes.

T4ΔHoc or T4ΔHocΔSoc used in engineered phage testing (Figure. 4-6) was propagated in *Escherichia coli* BL21(DE3). Bacteria were grown in LB medium supplemented with CaCl₂ and MgSO₄ to support phage infection; 200 mL cultures were grown at 37°C with shaking to mid-log phase. A 10 mL aliquot was infected with phage and incubated at 37°C for 10 min to allow adsorption, then returned to the remaining culture volume and incubated overnight at 37°C with shaking to allow phage amplification. Cultures were clarified by centrifugation (15,000 × g, 20 min, 4°C) and filtered through 0.22 µm filters to generate crude phage lysate.

### Bacteriophage purification by hydrophobic interaction chromatography (HIC)

For this work, we used established protocols^40^. In brief, clarified phage lysate was mixed 1:1 with high-salt binding buffer (potassium phosphate, pH 7.0) and purified by hydrophobic interaction chromatography on an ÄKTA purifier system equipped with a hydroxyl (OH) monolith column. Bound phage was eluted using a decreasing salt gradient; peak fractions were pooled and dialyzed into 1× PBS via two 2-hour dialysis steps followed by an overnight dialysis at 4°C, then filtered through 0.22 µm filters. Phage titers were measured post-propagation, post-purification, and post-dialysis to confirm recovery at each step.

### Diagnostic test for complement functions (WIESLAB)

Sera from healthy human volunteers (N = 8) were obtained as described above. WIESLAB Complement system Classical Pathway, WIESLAB Complement system Alternative Pathway and WIESLAB Complement system MBL Pathway kits (EuroDiagnostica, Sweden) were applied to assess the function of complement activity in human sera after incubation with phages. Ninety microliters of fresh human serum was incubated with 10 µL of PBS or highly purified T4 or T4ΔHoc phage preparations (final concentration 10^10^ PFU/mL) in 37°C for 10 min. Then, appropriate dilutions of all samples were added to pathway-specific plates and further steps were performed according to the manufacturer’s instructions. Briefly, plates were incubated at 37°C, for 1 h, washed 3 times with 300 µL of wash solution, and 100 µL of alkaline phosphatase-labelled antibodies to C5b-9 were added to wells. After 30 min incubation at room temperature, plates were again washed and 100 µL of substrate solution was added. Absorbance at 405 nm was measured after 30 min incubation at room temperature. Results were calculated as per cent of positive control (freezed dried human serum), which was delivered with the kits.

### Construction of peptide–Soc fusion constructs

Peptide sequences (a Compstatin variant; a Factor H–binding peptide; and a hydrophilic peptide) were synthesized by ANSA Biotechnology into a pET-Amp expression vector containing a ribosome binding site for expression in *E. coli* BL21(DE3). Each peptide was genetically fused to the C-terminus of the T4 small outer capsid (Soc) protein, a non-essential decoration protein that assembles on the phage capsid.

### Production of peptide-displaying bacteriophage

Expression plasmids encoding Soc–linker–peptide constructs were transformed into chemically competent BL21(DE3) cells and selected on LB agar containing ampicillin (100 µg/mL). Transformed colonies were grown overnight at 37°C with shaking (250 rpm), then diluted into 150 mL fresh LB with ampicillin and grown to mid-log phase (OD₆₀₀ ≈ 0.7). A 5 mL sample was collected prior to induction, and the remaining culture was pelleted (8,000 × g, 10 min, 4°C) and resuspended in fresh LB lacking antibiotics and supplemented with 0.2 mM IPTG to induce Soc–peptide fusion expression. Approximately 15 min after induction, a second 5 mL sample was collected for SDS-PAGE, and T4ΔHocΔSoc phage (lacking native Soc) was added to the culture at ∼10^6^ PFU/mL to initiate infection. Infected cultures were incubated at 37°C with shaking overnight, held at 4°C for ∼8 h, then clarified by centrifugation (15,000 × g, 20 min, 4°C) and filtered (0.22 µm) to obtain crude lysates, which were subsequently purified by HIC and dialyzed into 1× PBS as described above.

### SDS-PAGE analysis of Soc–peptide fusion protein expression

Bacterial pellets collected immediately prior to and 15 min following IPTG induction were resuspended in Laemmli loading buffer containing DTT, heated at 95°C for 5 min, then alkylated with iodoacetamide (500 mM) for 60 min in the dark. Samples were clarified by centrifugation (16,000 × g, 25 min) and resolved on 10–20% gradient polyacrylamide gels (100 V until entering the resolving gel, then 150 V). Gels were stained with Coomassie Brilliant Blue for ∼1 h, destained (methanol:acetic acid:water, 40:10:50) for ∼3 h, and rinsed prior to imaging.

### Anti-Soc ELISA for detection of peptide-displaying phage

Purified phage was diluted to 1 × 10^9^ PFU/mL in 1× PBS, and 100 µL was added per well of a 96-well plate, with parallel wells coated with either 1x PBS or heat-treated (80°C, 30 min) phage, in duplicate. Plates were incubated overnight at 4°C, washed, and blocked with SuperBlock buffer (1:5 in PBS) for 45 min. Primary detection used rabbit anti-Soc antibody (1:2000, 37°C, 2 h); secondary detection used HRP-conjugated anti-rabbit IgG (1:7500, room temperature, 1 h, dark). Signal was developed with TMB substrate (25 min, dark), stopped with 2 N H₂SO₄, and absorbance was read at 450 nm and 550 nm (reported as A_450_ − A_550_).

### Bacteriophage neutralization by human serum

Phage preparations were diluted in 1× PBS to 1 × 10^7^ PFU/mL. For each condition, 10 µL of phage was combined with 10 µL of active serum, heat-inactivated serum (56°C, 1 h), cobra venom factor (CVF)-treated serum, or 1× PBS as a control. For CVF treatment, 25 µL of serum was incubated with 1 µL CVF (250 U/mL) at 37°C for 1 h prior to use. Phage–serum mixtures were incubated at 37°C for 1 h, then serially diluted in PBS and plated by combining 50 µL of each dilution with 100 µL E. coli BL21(DE3) and 5 mL molten top agar (2%), overlaid onto LB agar plates. Plates were incubated overnight at 37°C, and plaques were counted the next day to calculate PFU/mL. Phage survival was expressed as log_10_(PFU_PBS/PFU_serum). This experiment was done in replicate across different days.

### Phage circulation in C3-deficient mice

Wild type C57Bl6/J (N = 6 or 8) and B6;129S4-*C3^tm1Crr^*/J (C3-deficient) (N = 6 or 8) male mice (Charles Rivers) were injected intraperitoneally (i.p.) with 10^9^ PFU/mouse of purified A3R or T4 phage preparations. Blood was collected from the tail vein into heparinized tubes after 2, 5, 10, 15 and 25 hours. Phage titers in each blood sample were determined using plaque assay.

### Phage circulation in C1-deficient mice

Wild type C57Bl6/J (N = 6) and B6N(Cg)-*C1qa**^tm1b(EUCOMM)Wtsi^***/3J (C1-deficient) (N = 6) male mice (Charles Rivers) were used, either naïve or pre-immunized with Hoc protein (25 µg/mouse s.c.) 5 days prior to the experiment. Animals were injected i.p. with 10^9^ PFU/mouse of purified T4 phage preparation. Blood was collected from the tail vein into heparinized tubes after 2, 5, 10 and 25 hours. Phage titers in each blood sample were determined using plaque assay.

### *In vivo* pharmacokinetics of peptide-displaying bacteriophages

Male C57BL/6 mice (6–8 weeks old) were immunized intraperitoneally (i.p.) with 25 µg of the T4 major capsid protein gp23 14 days prior to phage administration. Following immunization, mice received a single i.p. injection of 1 × 10⁹ PFU/mouse of either T4ΔHoc (n = 11), T4ΔHocΔSoc (n = 13), T4-HydroP (n = 13), or T4-FactorH.BP (n = 10). Whole blood was collected from individual mice at 1, 3, and 6 h following phage administration to assess circulating phage persistence. Blood samples were serially diluted in 1× PBS, and infectious phage titers were quantified by plaque assay using *Escherichia coli* BL21(DE3) as the bacterial host.

### Assessment of anti-phage antibody responses following gp23 immunization

To evaluate antibody responses following gp23 immunization, serum was collected on day 12 from six gp23-immunized mice and six PBS-injected control mice. ELISA plates were coated with 100 µL/well of purified T4ΔHoc phage (5 × 10^9^ PFU/mL) and incubated overnight at 4°C. Plates were washed with PBS and blocked with 1× SuperBlock buffer for 45 min at room temperature. Mouse serum was diluted 1:100 in PBS, and 100 µL was added per well and incubated at 37°C for 2 h. Following washing with PBS containing 0.05% Tween 20, bound antibodies were detected using HRP-conjugated goat anti-mouse IgG or IgM secondary antibodies (1:10,000) for 1 h at room temperature. Signal was developed using TMB substrate for 25 min in the dark and stopped with 2 N H_2_SO_4_. Absorbance was measured at 450 and 550 nm and reported as A_450_ − A_550_.

### Immunogold labeling of phage-associated C3 by transmission electron microscopy

A 5-µL aliquot of bacteriophage suspension (1 × 10^11^ PFU/mL) was applied to Formvar/carbon-coated nickel grids (400 mesh; Agar Scientific), and phage particles were allowed to adsorb for 5 min at room temperature in a humidified chamber. The grids were subsequently blocked with 1% (w/v) bovine serum albumin in PBS for 45 min at room temperature and washed six times with PBS for 1 min each. Grid-adsorbed bacteriophages were then exposed to healthy-donor human serum diluted 1:100 in PBS for 30 min at 37 °C, followed by six washes with PBS. Deposition of complement C3 fragments on phage particles was assessed by immunogold labeling using polyclonal rabbit anti-human C3c antibody (Dako; 1:200 in PBS) for 30 min at room temperature. Following six washes with PBS, the grids were incubated with a goat anti-rabbit IgG secondary antibody conjugated to 5-nm colloidal gold particles (Sigma; 1:20 in PBS) for 1 h at room temperature. Grids were then washed six times with PBS, negatively stained with 2% uranyl acetate, and examined using a JEM-1200EX transmission electron microscope (JEOL) operated at an accelerating voltage of 80 kV.

### Structural reconstruction of T4 capsid variants

Structural representations of the T4 capsid were generated in UCSF ChimeraX using the 3.3-Å cryo-EM structure of the bacteriophage T4 isometric capsid (PDB: 5VF3; EMDB: EMD-8661)^41^. The biological icosahedral assembly was displayed as a molecular surface, with the major capsid protein gp23, vertex protein gp24, and outer capsid proteins Soc and Hoc distinguished by color. T4ΔHoc and T4ΔHocΔSoc representations were generated from the same structural model by omitting Hoc for T4ΔHoc and both Hoc and Soc for T4ΔHocΔSoc, consistent with their respective capsid compositions. These representations are structure-based visualizations intended to illustrate differences in outer capsid protein composition and do not represent independently determined structures of T4ΔHoc or T4ΔHocΔSoc.

### Statistical analysis

Statistical analyses were performed using GraphPad Prism. Depending on the experimental design, data were analyzed using one-way or two-way ANOVA, mixed-effects models (REML), or unpaired Welch’s t-tests, followed by Tukey’s, Dunnett’s, or Šídák’s multiple comparisons tests where appropriate, as indicated in the respective figure legends. Statistical significance was defined as P < 0.05.

## Author contributions

M.A., P.L.B., K.D., P. M., conceived the study. M.A., K.D., P.M. designed the experiments. M.A., B. R., P. M., Z.K., I.R., A.K., K.D., T.L. performed the experiments. P.L.B., K.D., contributed reagents. M.A., I.R., P.M. propagated and purified phages. M.A., P.L.B, K.D., P.M. analyzed and interpreted the data. M.A., P.L.B., K.D., P.M. wrote the manuscript. All authors reviewed the manuscript.

## Funding Acknowledgements

P.L.B discloses support from the Cystic Fibrosis Foundation CHEN21F0; Cystic Fibrosis Research Institute; Stanford Woods Institute for the Environment, National Institutes of Health grants P01AI1960471, K24 AI166718, R01 AI182349, R21AI194044, and R01 EB038154. KD and PM acknowledge support from the National Science Centre in Poland grant UMO-2021/40/Q/NZ7/00202. The contents are those of the authors and do not necessarily represent the views of the funding agencies.

## Data availability statement

All data supporting the results in this study are available within the paper and its Supplementary Information. Additional data are available from the corresponding author on reasonable request.

