## Supplemental Data 1 for "Engineering Bacteriophages to Overcome Complement-Mediated Neutralization"

**SUPPLEMENTARY FIGURES**

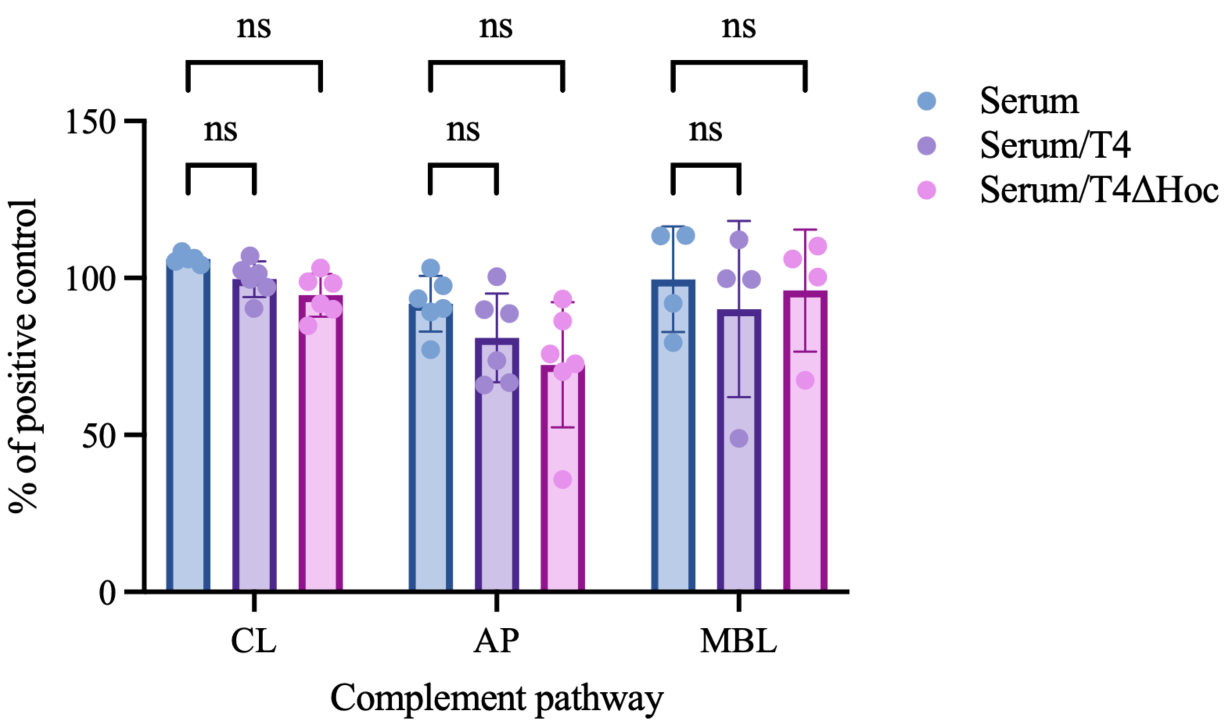


**Supplementary Figure 1. Effects of T4 and T4ΔHoc on complement pathway activity.** Classical (CL), alternative (AP), and lectin (MBL) complement pathway activity was assessed using WIESLAB® Complement System assays following incubation of human serum with PBS, T4, or T4ΔHoc ($10^{10}$ PFU/mL). Activity is expressed as a percentage of the positive control. Points represent individual donor measurements (up to n = 6 donors per condition), and bars represent mean ± SD. Statistical comparisons were performed using a two-way mixed-effects model (REML), with measurements matched by donor across complement pathways and phage treatments, followed by Dunnett's multiple-comparisons test comparing each phage-treated condition with untreated serum within each pathway. No individual comparisons were statistically significant following adjustment (all adjusted $P > 0.05$).

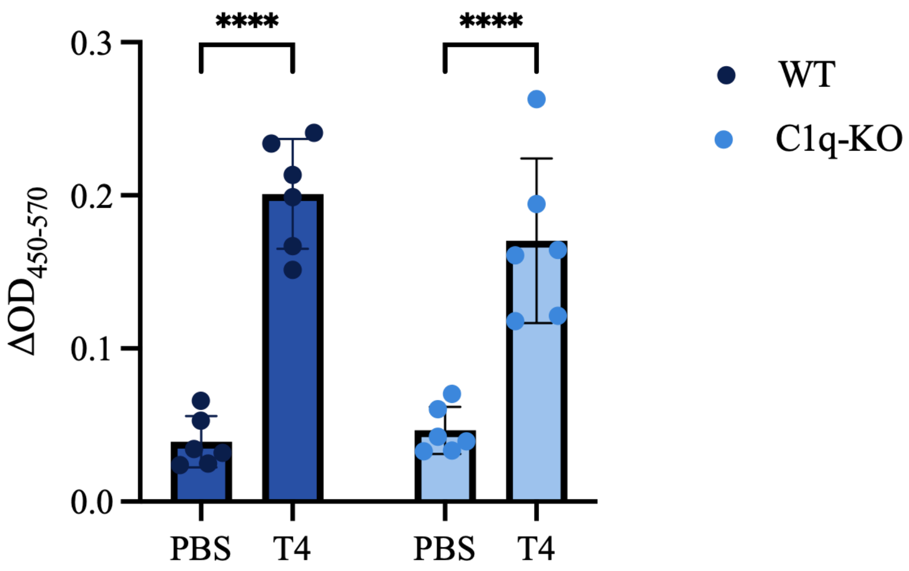


**Supplementary Figure 2. Anti-T4 antibody responses in Hoc-immunized WT and C1q-KO.** Serum antibody reactivity to T4 phage was assessed by ELISA in WT and C1q-KO mice immunized with the T4 outer capsid protein Hoc (n = 6 per genotype). T4-coated wells were compared with PBS controls. Data are presented as mean ± SD, with individual mice represented by dots. Statistical significance was determined by ordinary two-way ANOVA followed by Šídák's multiple comparisons test comparing T4 and PBS within each genotype (****$P < 0.0001$).

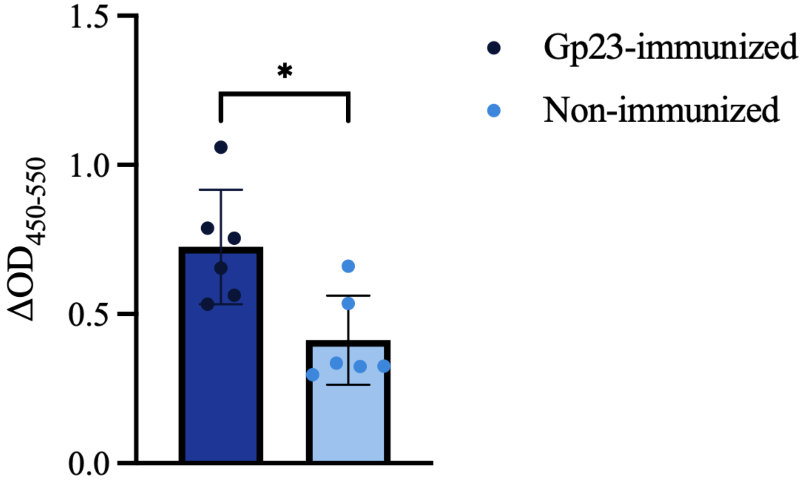


**Supplementary Figure 3. Gp23 immunization increases antibody reactivity against T4ΔHoc.** Serum antibody reactivity to T4ΔHoc phage was assessed by ELISA in mice immunized with the T4 major capsid protein Gp23 or non-immunized controls (n = 6 per group). Antibody reactivity was measured as $\Delta OD_{450-550}$. Data are presented as mean ± SD, with individual mice represented by dots. Statistical significance was determined using a two-tailed unpaired Welch's t-test (*P = 0.0112).
